# Metabolic reprogramming and stress mitigation of *Chlamydomonas reinhardtii* using protective metal-phenolic networks

**DOI:** 10.64898/2026.03.11.711231

**Authors:** Wenting Liao, Chenyu Wang, Bohan Cheng, Joseph J. Richardson, Kanjiro Miyata, Hirotaka Ejima

## Abstract

Single-cell encapsulation is widely used in biotechnology to protect highly sensitive living cells from environmental stress. However, the impact of encapsulation on biological function and metabolism remains rarely explored despite its importance for understanding stress response and modulating the production of valuable metabolites. Herein, we show that coating individual cells with protective metal-phenolic networks not only improves survival against stress, but also modifies cellular metabolism. Importantly, this encapsulation technique induces a reversible state of quiescence, which enables accumulation of high-energy-density compounds through selective nutrient diffusion and the resulting carbon flux redistribution. Specifically, light exposure favors the accumulation of nearly eightfold higher starches, while incubation in darkness leads to twofold higher lipid accumulation compared to native cells. This metabolic engineering approach via individual cell encapsulation expands the cell editing toolbox and will facilitate applications in synthetic biology, bioengineering, and cell-based therapeutics.

**Graphical Abstract:** 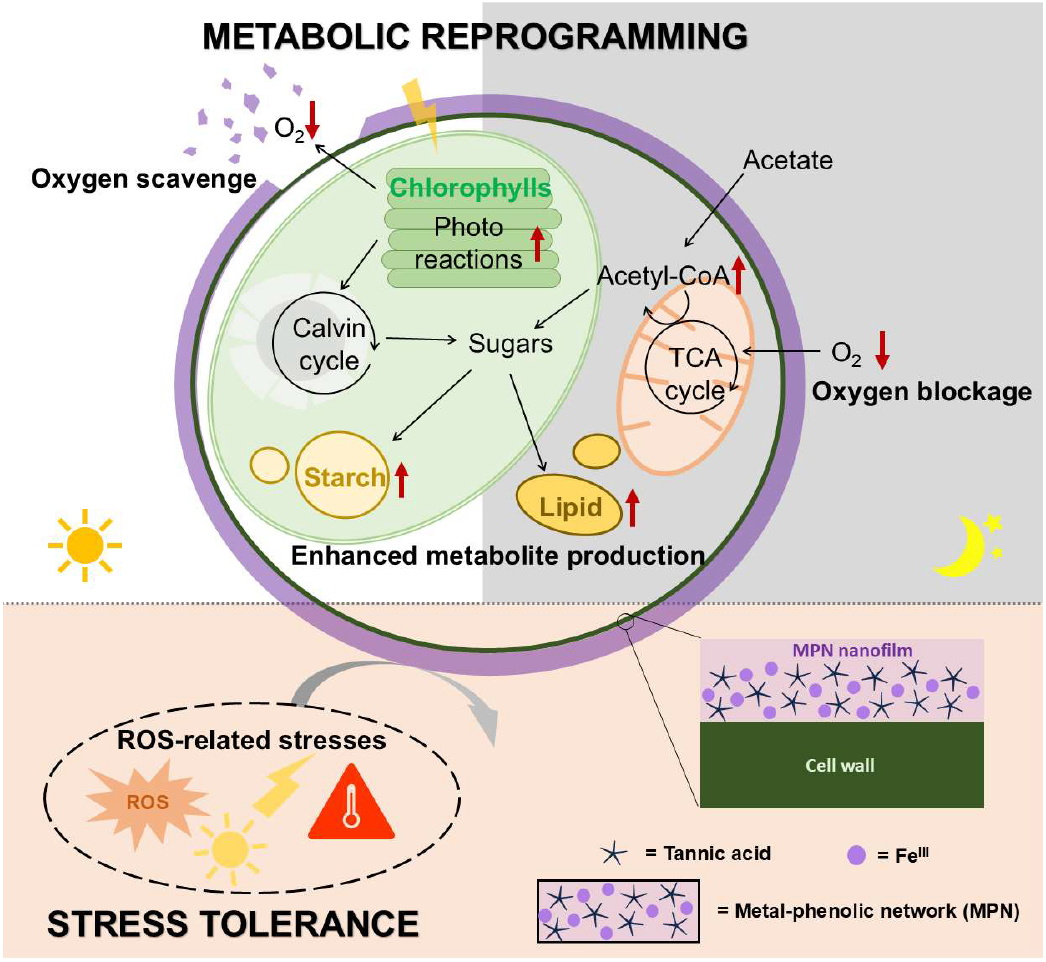

## Introduction

The encapsulation of living cells within artificial shells has recently emerged as an effective strategy to protect fragile cells against environmental stress.^1^ Among the various protective encapsulating technologies studied to date, metal-phenolic networks (MPNs) are particularly promising materials due to their facile and benign assembly from biocompatible building blocks, and their tunable permeability and intrinsic antioxidant properties.^2,3^ Importantly, the reversibility of metal-phenolic coordination allows for dynamic control over the stability of the MPN shell,^4,5^ meaning that the cells can be selectively encapsulated and released in response to environmental triggers. However, MPN-based cell encapsulation has primarily focused on preserving cell viability,^6,7^ and are often regarded as rigid protective layers that passively preserve cellular integrity,^8,9^ rather than active coatings controlling cell function on a deeper level.

Cells respond to a wide range of cues in their surrounding environment, and to date metabolic engineering has focused on either changing external cues in growth conditions (i.e., by adding signaling molecules^10^ or changing temperature^11^) or reprogramming the cells at the genetic level (i.e., genetic engineering^12^). MPNs offer a unique opportunity to control the external signals reaching encapsulated cells due to their antioxidant activity^13^ and tunable permeability^14^, which could potentially influence the intracellular redox balance and energy metabolism of cells. For example, MPN encapsulation can change the microenvironment and enable methane production of strictly anaerobic methanogens at atmospheric oxygen levels.^13^

In this study, we selected *Chlamydomonas reinhardtii* (*C. reinhardtii*) as the model organism for a proof-of-concept investigation into how MPN encapsulation can modulate cellular metabolism under different growth conditions. As a photosynthetic microorganism exhibiting distinct metabolic modes under light and dark conditions, *C. reinhardtii* provides a unique platform to explore MPN encapsulation, and we found that MPNs not only protect the cells from various stress triggers, but also modulate oxygen diffusion into the cells, which promotes photo reactions by scavenging the produced oxygen under light and constrains the tricarboxylic acid (TCA) cycle in darkness, thereby redirecting carbon flux toward fatty acid and triacylglycerol (TAG) accumulation. Specifically, incubation under light triggered the disassembly of MPN coatings together with an increased accumulation of starches, while if the incubation was performed in darkness, lipids accumulated at an increased level over 7 days. Collectively, these results highlight that cell encapsulation not only impacts cell division and viability, but is also a unique approach to modify metabolism and the production of valuable metabolites.

## Results and Discussion

### Encapsulation of *C. reinhardtii* cells in MPN shells

To mimic the natural sporulation process, *C. reinhardtii* was encapsulated into tannic acid (TA)-iron MPN nanofilms (algae@MPN, Fig. 1a) with different layer numbers (algae@MPN×*n*) similar to our previous reports (Fig. S1).^4,15^ To confirm encapsulation, TA was labeled with DyLight 405 *N*-hydroxysuccinimide (NHS) ester-conjugated bovine serum albumin (BSA-DyLight405), and after MPN formation confocal microscopy images clearly showed that individual algae were encapsulated (Fig. 1b, Fig. S2). Specifically, the ring-shaped DyLight405-labeled MPN layers were blue, while the microalgae chlorophyll produced red autofluorescence in the cores and the merged image exhibited conventional core-in-shell structures confirming encapsulation. Moreover, the cell suspensions darkened as the number of MPN coating layers increased from 1 to 4 due to the strong visible light absorbance of the iron coordination center (Fig. 1c). Transmission electron microscopy further confirmed the formation of rough-surfaced MPN layers around the cells (Fig. 1d, Fig. S3). Similar to previous reports,^8^ the encapsulation process did not impair the viability of the encapsulated cells (Fig. 1e, Fig. S4). Moreover, the MPN shell was stable in cell culture media for at least 4 days when monitored on UV-inactivated cells (Fig. S5).

**Figure 1.**
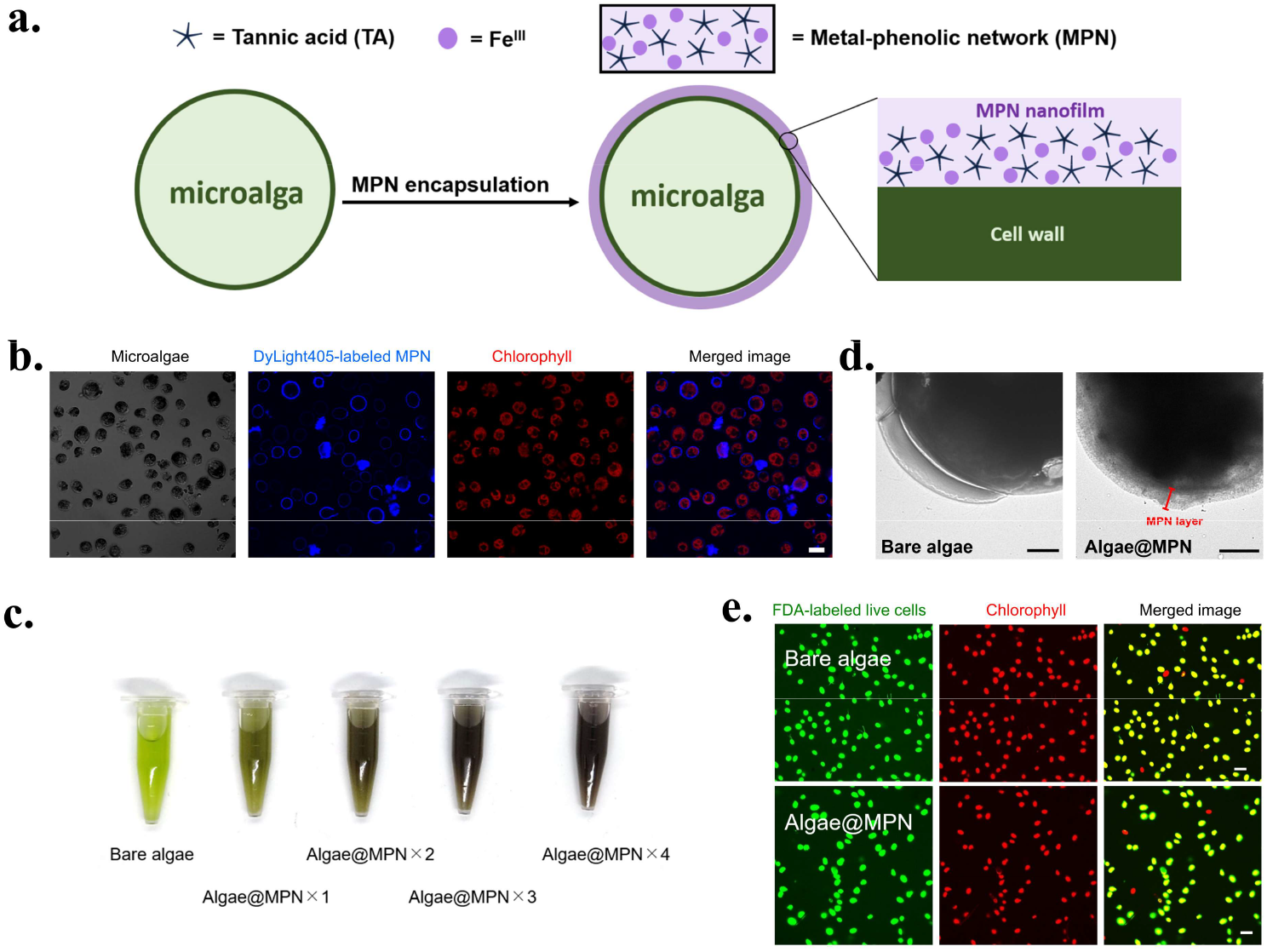
Encapsulation of microalgae, *Chlamydomonas reinhardtii* (*C. reinhardtii*), in metal-phenolic networks (MPNs). (a) Schematic illustration of the artificial sporulation process of *C. reinhardtii* using MPN shells. (b) Confocal laser scanning microscopy (CLSM) images of bare algae and algae@MPN using DyLight405-labeled MPN confirming encapsulation. (c) Photographs of suspensions of bare algae and MPN-encapsulated algae (algae@MPN). (d) Transmission electron microscopy images of bare algae and algae@MPN. (e) CLSM images of bare algae and algae@MPN using fluorescein diacetate (FDA, green) to confirm viability. In (b,c,d,e), independent experiments were performed (*n* = 3) with similar results. Scale bars: 10 μm (b,e); 2 μm (d).

### MPN encapsulation offers oxidative and environmental resilience

Previous reports have demonstrated that MPN encapsulation offers cytoprotection against various stressor, including toxic chemicals, ultraviolet (UV) radiation, heat, and nanoparticles.^8,16,17^ We confirmed that algae@MPN exhibited markedly enhanced viability under multiple reactive oxygen species (ROS)-related stresses, including oxidative challenge, UV irradiation, and heat shock, whereas bare algae showed pronounced susceptibility to all three (Fig. 2a–d). However, the changes to metabolism and biological function following encapsulation and exposure to stressors have never been studied to date, and therefore we employed mitochondrial membrane potential (MMP) assay to investigate the impact of oxidative stresses on mitochondria function. The measurement of MMP was conducted with JC-1 dye,^18^ which accumulates in the mitochondria, and emits red fluorescence at low membrane potentials and green fluorescence at high membrane potentials enabling quantification of mitochondrion health (Fig. S6–7).^19^ Interestingly, after exposure to oxidative stress (1 mM hydrogen peroxide, H_2_O_2_, 24 h), the mitochondrial function of algae@MPN was nearly identical to untreated algae, however when bare algae was exposed to the same oxidative stress they exhibited a significant decrease in mitochondrial health (Fig. 2e).

**Figure 2.**
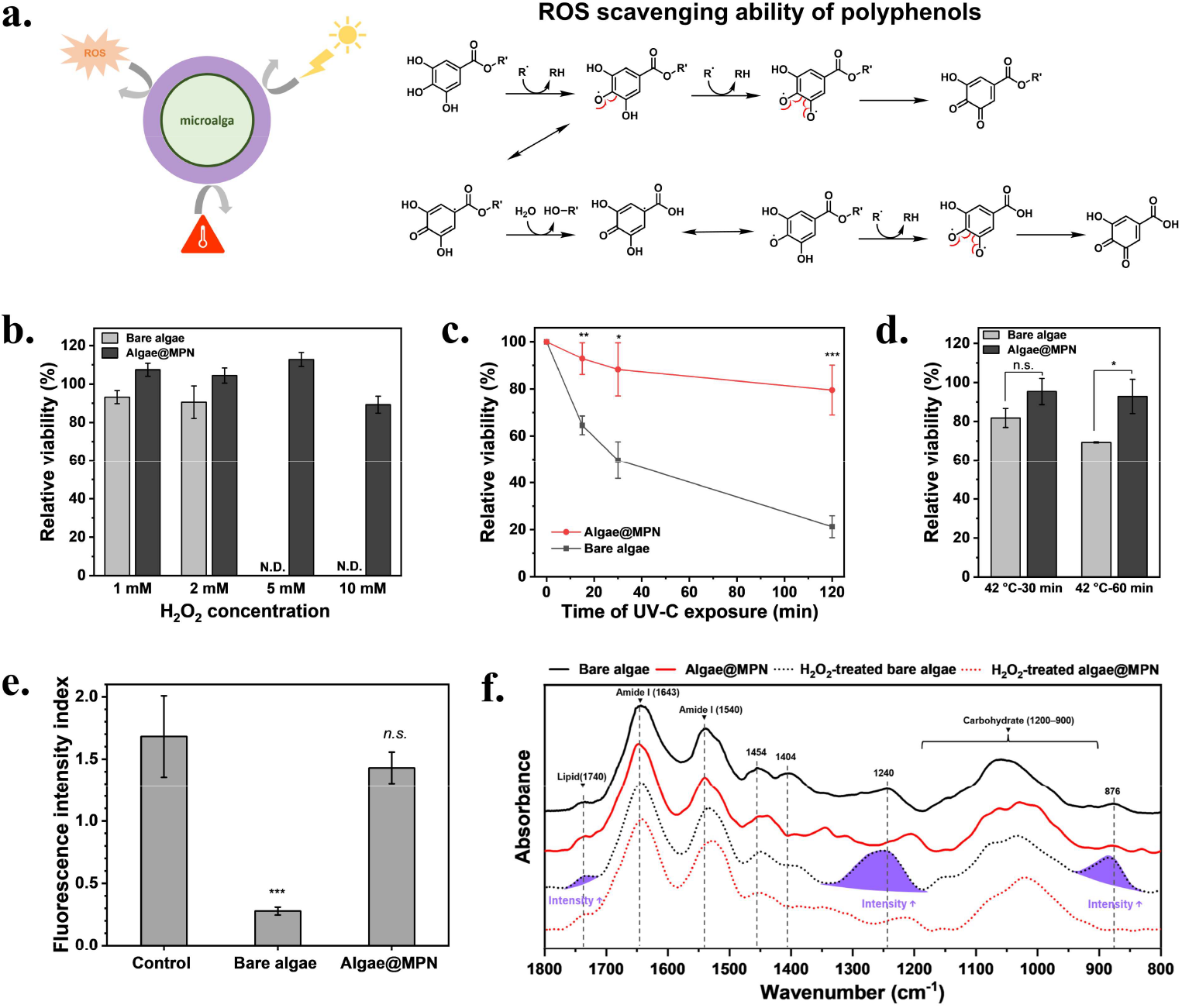
Enhanced environmental resilience of *Chlamydomonas reinhardtii* (*C. reinhardtii*) offered by metal-phenolic network (MPN) encapsulation. (a) Schematic illustration of protection of *C. reinhardtii* toward environmental stresses by antioxidant MPN shells. (b–d) Relative viability of bare algae and algae@MPN under oxidative stress, ultraviolet-C (UV-C) irradiation, and heat shock, respectively. (e) Mitochondrial membrane potential of untreated bare algae (Control), hydrogen peroxide (H_2_O_2_)-treated bare algae, and H_2_O_2_-treated algae@MPN, evaluated by JC-1 fluorescence intensity ratio. (f) Attenuated total reflectance Fourier transform infrared spectra of bare algae and algae@MPN before and after H_2_O_2_ treatment. Error bars represent standard deviations (*n* = 3). Statistical analysis was performed using one-way *ANOVA* (\*\*\**p* < 0.001, \*\**p* < 0.01, \**p* < 0.05, n. s. *p* ≥ 0.05). In (f), independent experiments were performed (*n* = 3) with similar results.

Attenuated total reflectance Fourier transform infrared (ATR-FTIR) spectroscopy further confirmed that the MPN shells were protecting against oxidative stress, as bare algae exhibited obvious changes in the 1735 cm^−1^, 1250 cm^−1^, and 882 cm^−1^ bands after H_2_O_2_ treatment, while the algae@MPN spectra resembled that of the pristine algae (Fig. 2f). Specifically, common peaks related to the major biomolecules in *C. reinhardtii* were observed in the ATR-FTIR spectra of all samples (Fig. 2f, and Table S1), and characteristic peaks of the functional groups in TA were observed in the algae@MPN spectra (Fig. S8, and Table S2). We further quantified the changes in specific biomolecules using the amide I band at 1643 cm^−1^ as the reference (Table S3) and saw that oxidative stress caused lipid peroxidation, potential cleavage of nucleic acids, and structure alteration of the sugar-rich glycoprotein cell wall. However, these changes were greatly reduced when the microalgae were protected with MPN shells. Overall, MPN encapsulation significantly enhanced the oxidative resilience of cells, and because oxygen and its radical derivatives are important metabolites of various cellular activities, we assumed that MPN encapsulation might also influence metabolic processes.

### Short-term encapsulation-induced metabolic shift

To investigate the influence of MPN encapsulation on metabolism, we first focused on the respiration process, and the microalgae were cultured in darkness to deactivate photosynthesis. MMP increased immediately after encapsulation, suggesting that the MPN shell may transiently limit oxygen diffusion (Fig. 3a). This limited oxygen diffusion is likely due to chemical bonding of oxygen to the metal coordination center of the MPN. Reduced oxygen availability could impede the final steps of the TCA cycle where oxygen serves as the final electron acceptor to produce adenosine triphosphate (ATP), causing electron accumulation in mitochondria and a temporary rise in MMP. With longer incubation in the dark (up to 24 h), MMP gradually declined as cellular metabolism slowed. Notably, algae@MPN showed a faster decrease in MMP than bare algae, likely due to stronger oxygen limitation reducing ATP production and further suppressing metabolism. Even so, the lowest measured JC-1 red-to-green fluorescence ratio (Fig. S9, 3.5 ± 0.4) remained within the range typically associated with highly energized mitochondria,^20^ indicating that although MPN encapsulation limited oxygen availability, it only induced mild stress and did not cause irreversible damage.

**Figure 3.**
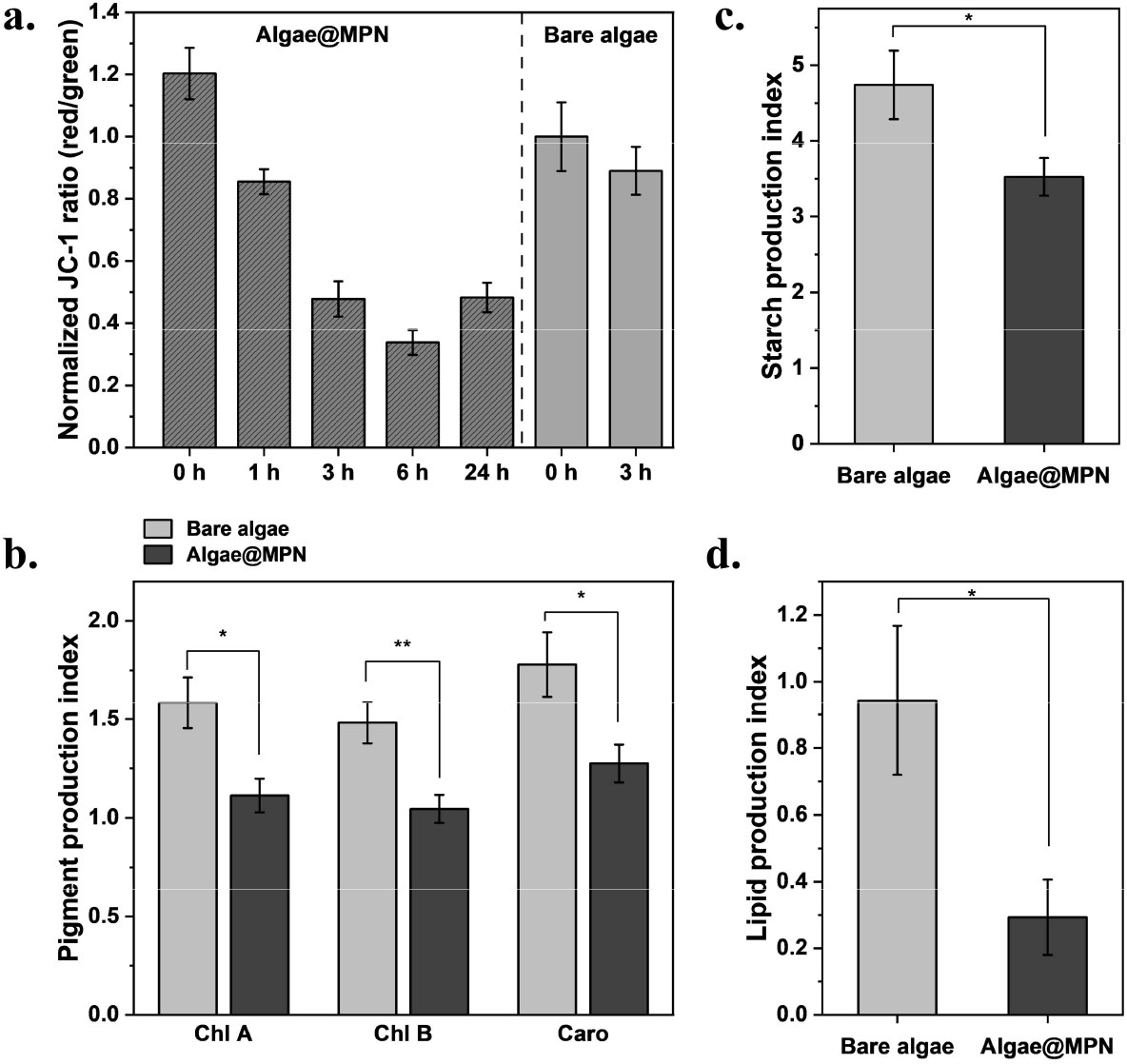
Mild metabolic adjustment of *Chlamydomonas reinhardtii* in the early phase after metal-phenolic network (MPN) encapsulation. (a) Normalized red-to-green fluorescence ratio of algae@MPN and bare algae measured by the JC-1 mitochondrial membrane potential (MMP) assay after incubation in darkness. (b) Pigment production index of bare algae and algae@MPN after 1-day of incubation in the dark. Chl A: chlorophyll a; Chl B: chlorophyll b; Caro: full carotenoids. (c,d) Starch and lipid production indices of bare algae and algae@MPN after 1-day of incubation in the dark. Error bars represent standard deviations (*n* = 3). Statistical analysis was performed using one-way *ANOVA* (\*\*\**p* < 0.001, \*\**p* < 0.01, \**p* < 0.05, n. s. *p* ≥ 0.05).

To further examine the metabolic slowdown caused by MPN encapsulation, we evaluated the production of three key metabolites (pigment, starch, and lipid) in *C. reinhardtii* after 1 day of dark cultivation. To account for variability in the initial states of independent cell groups, we introduced a metabolite production index, calculated by normalizing metabolite levels to their pre-treatment values, enabling more accurate comparisons across conditions. As expected, algae@MPN accumulated fewer metabolites than bare algae (Fig. 3b–d). Interestingly, the lipid index of bare algae was close to 1, indicating minimal utilization of stored energy in darkness, while the lipid index of algae@MPN dropped to 0.3, suggesting that encapsulated cells consumed much of their lipid reserve to cope with dark conditions. Overall, this reduced metabolite production reflected a transition to a protective, low-activity, quiescent-like state in the early phase after MPN encapsulation.

### Long-term encapsulation-induced metabolic shift

Because MPN encapsulation restricts oxygen, we hypothesized that prolonged dark cultivation might further alter the metabolism of *C. reinhardtii*, particularly in energy-source accumulation, similar to nutrient-deprivation-induced lipid enhancement (e.g., nitrogen or phosphorus starvation).^21,22^ Nile red staining was used to visualize intracellular lipids, and CLSM images showed substantial lipid accumulation in algae@MPN after 4 days in darkness (Fig. 4a). Interestingly, bare algae produced almost no lipids after 4 days, however both groups exhibited lipid accumulation after 7 days (Fig. S10). To quantify this change, the total lipids were extracted using the Bligh and Dyer method^23^ from cultures seeded at 1×10^5^ cells/mL and grown in continuous darkness for 7 days. Algae@MPN×4 accumulated ∼26% more lipids than bare algae, though this difference was not statistically significant (Fig. 4b). However, increasing the shell thickness (algae@MPN×8) resulted in significantly higher lipid accumulation, likely due to improved oxygen blocking and longer retention of the encapsulation layer.

**Figure 4.**
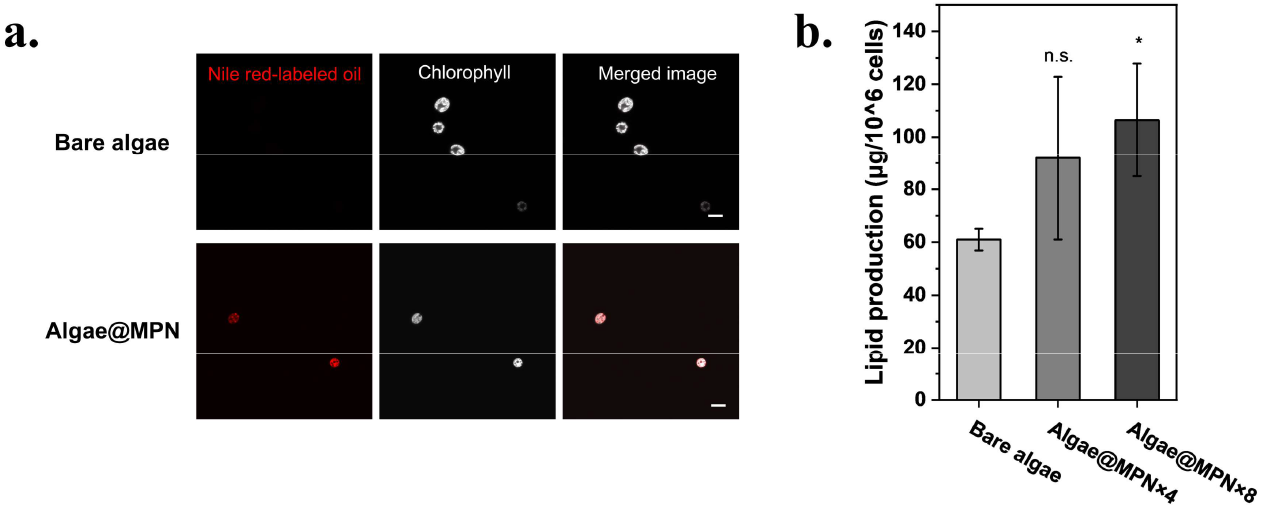
Enhanced lipid accumulation of *Chlamydomonas reinhardtii* by MPN encapsulation under dark cultivation. (a) Confocal laser scanning microscopy (CLSM) images of Nile red-stained bare algae and algae@MPN after dark cultivation for 4 days. Lipid droplets (left) and chlorophyll autofluorescence (middle) are shown, together with merged images (right). (b) Lipid production of bare algae and algae@MPN subjected to multiple encapsulation cycles (×4 and ×8) after dark cultivation. In (a), independent experiments were performed (*n* = 3) with similar results. Scale bars: 10 μm. Error bars represent standard deviations (*n* = 3). Statistical analysis was performed using one-way *ANOVA* (\**p* < 0.05, n. s. *p* ≥ 0.05).

### Light-triggered disassembly of MPN layers and metabolic shift following release

As we reported previously,^4^ ROS can trigger MPN disassembly. Because *C. reinhardtii* generates singlet oxygen and related radicals during photosynthesis, we tested if light stimulation could induce self-disassembly of the MPN layer from algae@MPN. While most MPN layers remained on the cell surface after 24 h in darkness, exposure to light for 24 h caused disassembly (Fig. 5a), indicating the responsiveness of MPN encapsulation towards the cellular metabolism under light conditions. Moreover, algae@MPN maintained photosynthetic pigment profiles comparable to bare algae under light (Fig. 5b), suggesting that light-induced MPN disassembly allowed recovery of normal photosynthetic function. Specifically, the pigment production indices after 24 h of light exposure showed minimal differences in chlorophyll and carotenoid levels, and their net production over longer light exposure (7 days) was also similar (Fig. S11).

**Figure 5.**
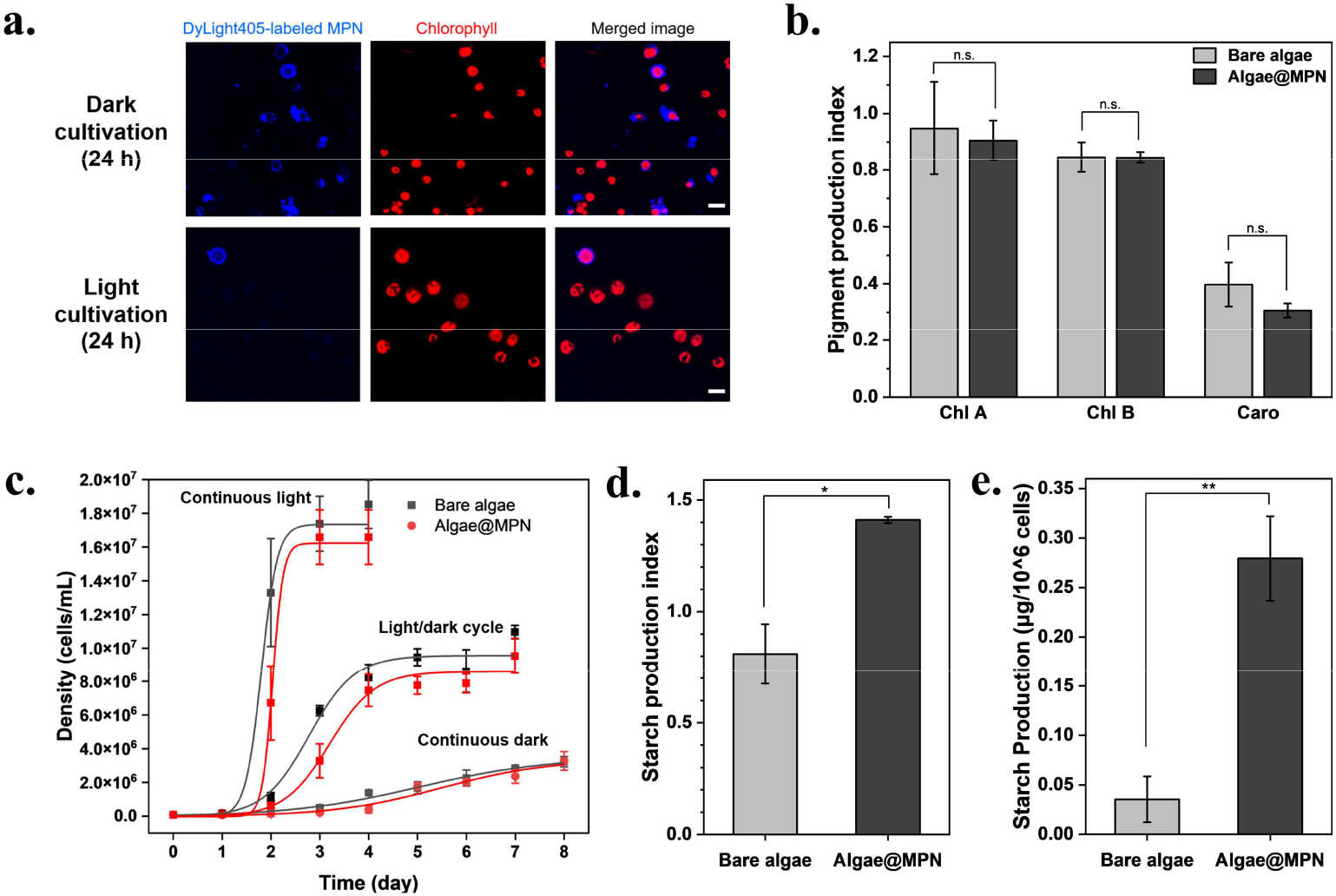
Light-triggered disassembly of metal-phenolic network (MPN) layers and associated metabolic shift of *Chlamydomonas reinhardtii*. (a) Confocal laser scanning microscopy images of DyLight405-labeled bare algae and algae@MPN after incubation under light or in darkness. DyLight405-labeled MPN layer (left) and chlorophyll autofluorescence (middle) are shown, together with merged images (right). (b) Pigment production indices of bare algae and algae@MPN under 24 h light exposure, including the production of chlorophyll a (Chl A), chlorophyll b (Chl B), and total carotenoids (Caro). (c) Cell growth profiles of bare algae and algae@MPN cultured under different light conditions. (d,e) Starch production of bare algae and algae@MPN under light, evaluated by short-term (24 h) production indices and long-term (7 days) net production. In (a), independent experiments were performed (*n* = 3) with similar results. Scale bars: 10 μm. Error bars represent standard deviations (*n* = 3). Statistical analysis was performed using one-way *ANOVA* (\*\**p* < 0.01, \**p* < 0.05, n. s. *p* ≥ 0.05).

Growth kinetics after disassembly further demonstrated the light-responsive nature of MPN encapsulation. Under continuous light, light/dark cycles, or continuous darkness, both bare algae and algae@MPN showed typical sigmoid growth, with encapsulated cells exhibiting longer lag phases and negligible growth (Fig. 5c, and Table S4). Notably, longer light exposure increased growth, supporting the idea that oxygen and ROS generated during photosynthesis gradually disassemble the MPN shells allowing for proper cell division. Algae@MPN also displayed growth rates slightly higher than bare algae following release (Table S4). After 1 day of light cultivation, the starch levels increased significantly in algae@MPN but were partially consumed in bare algae (Fig. 5d). This suggested that during the encapsulation-induced lag period, cells continued accumulating internal metabolites (e.g., starch) but were physically restricted from division, and once the MPN shell was disassembled these stored resources enabled faster growth. Even after prolonged light exposure (7 days), when MPN layers were largely disassembled, algae@MPN still showed higher net starch production (Fig. 5e), which might result from MPN residues scavenging excess ROS and oxygen, reducing photodamage and enhancing photosynthetic efficiency, thereby promoting carbon dioxide (CO_2_) fixation and starch accumulation.

### Conceptual model of encapsulation-induced metabolic reprogramming

Our results collectively showed that MPN encapsulation triggers metabolic reprogramming in *C. reinhardtii* rather than functioning merely as a passive protective layer. Based on changes in pigment levels, starch and lipid accumulation, and stress tolerance, we developed a mechanistic model describing how MPN coatings regulate both photosynthesis and respiration (Scheme 1).

Under light, photosynthetic activity remains largely intact, as indicated by comparable pigment content between bare and coated cells, suggesting that the MPN layer does not impair light harvesting. Instead, its antioxidant properties and selective gas permeability subtly modify the cellular microenvironment. While CO_2_ and acetate can diffuse normally, oxygen diffusion is partially restricted, slightly shifting the redox balance and causing transient accumulation of equivalent amounts of reducing molecules (e.g., NADPH). This favors cyclic electron flow in photosystem I (PSI), providing additional ATP to support biosynthesis. The Calvin cycle continues CO_2_ fixation, and the excess photosynthate is stored as starch, consistent with the elevated starch levels observed in encapsulated cells. Accordant with this mechanism, encapsulated cells exhibited significantly greater net starch accumulation under prolonged light exposure, even though the MPN layer is already partially disassembled.

When photosynthesis ceases in darkness, coated cells also exhibit altered respiratory behavior. Immediately after encapsulation, limited oxygen availability increases MMP due to impaired terminal electron transport, marking a temporary low-activity, protective state. MMP gradually normalizes within 24 h, but pigment, starch, and lipid contents remain lower than in bare cells, indicating a transient suppression of biosynthesis during microenvironmental adaptation and a shift toward metabolic quiescence.^24^ After this phase, acetate is metabolized into acetyl-coenzyme A (acetyl-CoA), supporting fatty acid and TAG synthesis instead of the TCA cycle as oxygen, the final electron acceptor of the TCA cycle, is limited. Overall, MPN encapsulation acts as a dynamic biochemical interface that modulates the redox balance and energy allocation through antioxidant buffering and partial gas regulation. This dual functionality not only enhances stress tolerance but also redirects metabolic fluxes toward energy-dense storage compounds, offering a generalizable strategy to tune cell metabolism through engineered material–cell interfaces.

**Scheme 1.**
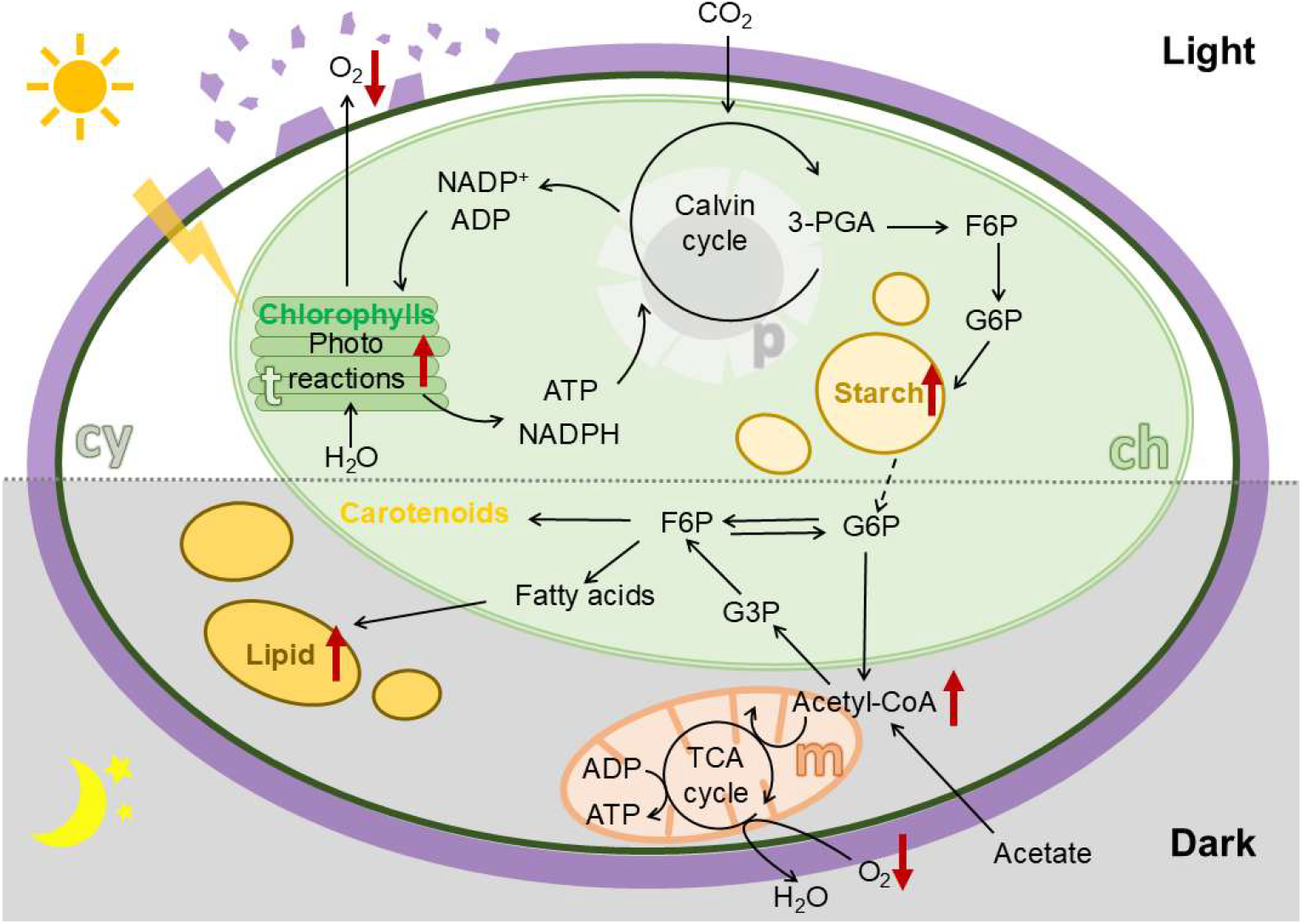
Schematic illustration of the cellular metabolisms of algae@MPN. t: thylakoid, ch: chloroplast, cy: cytosol, p: pyrenoid, m: mitochondria, 3-PGA: 3-phosphoglycerate, F6P: fructose 6-phosphate, G6P: glucose 6-phosphate, acetyl-CoA: acetyl coenzyme A.

## Conclusion

In summary, we demonstrated that coating individual *C. reinhardtii* cells with MPNs leads to a reversible protective layer that enhances survival against oxidative stress, UV radiation, and temperature fluctuations. Beyond protection, the coating actively modulates cellular metabolism into a reversible state of quiescence: in darkness, it transiently suppresses biosynthetic activity but subsequently promotes lipid accumulation, whereas under light, it enhances starch storage. The coating gradually disassembles under light, allowing cells to regain their native surface properties and cell division in a controlled manner. Together, these results establish MPNs as functional and reversible cell coatings for metabolic regulation, offering a versatile platform for synthetic biology, bioengineering, and cell-based technologies requiring robust yet adaptable living systems.

## Supporting Information

The authors have cited additional references within the Supporting Information.^23,25^

## Acknowledgment

This research was partially supported by Japan Society for the Promotion of Science (JSPS) KAKENHI (Grant No. 20K20641 and 25K01824) and Japan Science and Technology Agency (JST) SPRING, Grant Number JPMJSP2108. This work was also supported by “Advanced Research Infrastructure for Materials and Nanotechnology in Japan (ARIM)” of the Ministry of Education, Culture, Sports, Science and Technology (MEXT), Grant Number JPMXP1224UT0117.

## Author contributions

Conceptualization, W.L. and H.E.; methodology, W.L., B.C., J.J.R., and H.E.; Investigation, W.L. and C.W.; writing—original draft, W.L.; writing—review & editing, W.L., C.W, B.C., J.J.R., K.M., and H.E.; funding acquisition, W.L. and H.E.; resources, M.K. and H.E.; supervision, H.E.

