## Supporting information for "Metabolic reprogramming and stress mitigation of *Chlamydomonas reinhardtii* using protective metal-phenolic networks"

### 1. Experimental Procedures

#### 1.1 Materials

Tannic acid (TA), iron (III) chloride hexahydrate ($FeCl_3 \cdot 6H_2O$), bovine serum albumin (BSA), agar, ammonium chloride ($NH_4Cl$), magnesium sulfate heptahydrate ($MgSO_4 \cdot 7H_2O$), calcium chloride dihydrate ($CaCl_2 \cdot 2H_2O$), dipotassium hydrogen phosphate ($K_2HPO_4$), potassium dihydrogen phosphate ($KH_2PO_4$), zinc sulfate heptahydrate ($ZnSO_4 \cdot 7H_2O$), boric acid ($H_3BO_3$), manganese chloride tetrahydrate ($MnCl_2 \cdot 4H_2O$), cobalt chloride hexahydrate ($CoCl_2 \cdot 6H_2O$), copper sulfate pentahydrate ($CuSO_4 \cdot 5H_2O$), ammonium heptamolybdate tetrahydrate ($(NH_4)_6Mo_7O_{24} \cdot 4H_2O$), ferrous sulfate heptahydrate ($FeSO_4 \cdot 7H_2O$), potassium hydroxide (KOH), dimethylformamide (DMF), 10× phosphate-buffered saline (PBS), fluorescein diacetate (FDA), acetone, propidium iodide (PI), hydrogen peroxide ($H_2O_2$), dimethyl sulfoxide (DMSO), ethanol, Nile red, methanol, chloroform, and sodium acetate were purchased from FUJIFILM Wako Pure Chemical Corporation (Osaka, Japan). JC-1 MitoMP detection kit was purchased from Dojindo Laboratories Co., Ltd (Kumamoto, Japan). Ethylenedinitrilotetraacetic acid disodium salt dihydrate ($Na_2 \cdot EDTA$) were purchased from Tokyo Chemical Industry Co., Ltd (Tokyo, Japan). DyLight™ 405 NHS ester was purchased from Thermo Fisher Scientific (MA, USA). Starch assay kit (SA20) was purchased from Sigma-Aldrich (MA, USA). Deionized (DI) water from a water purification system (Direct-Q UV3 Remote, Merck) was used. The recipe for the tris-acetate-phosphate (TAP) medium was described in Table S5–8. Before use, TAP media were sterilized by autoclave (LSX-500, Tomy Seiko,Tokyo, Japan), and all stock solutions are stored unsterilized at 4 °C.

#### 1.2 Culture of *Chlamydomonas reinhardtii*

*Chlamydomonas reinhardtii* (*C. reinhardtii*) was seeded and cultured on a TAP agar plate under natural sunlight at 25 °C. One single colony was picked from the agar plate and seeded in a 40 mL TAP liquid medium. The cells were cultured in an LED-equipped shaking incubator (LC-LED450W and BR-43FL·MR, Taitec, Saitama, Japan) at 25 °C at 120 rpm under continuous light. The light intensity was set to 60% of the maximum, where the maximum photon flux density was approximately 215 μmol photons $m^{-2}$ $s^{-1}$.

Oxidative stress was induced by adding a 10 μL $H_2O_2$ solution into a 1 mL cell suspension ($2.5 \times 10^6$ cells/mL), resulting in desired $H_2O_2$ concentrations. The cells were cultured under natural sunlight at 25 °C for 24 h. Ultraviolet (UV) stress was induced by exposing cell suspension ($2.5 \times 10^6$ cells/mL) to 254 nm UV-C light at an intensity of approximately 50 $\mu W/cm^2$ for a predetermined time. Thermal stress was induced by incubating cell suspensions ($2.5 \times 10^6$ cells/mL) at 42 °C for a predetermined time.

#### 1.3 Formation of microalgal spores

The *C. reinhardtii* microalgae cells were collected by centrifugation (2000 × g, 5 min), washed twice with fresh TAP media, and resuspended in TAP media with their density adjusted to $1 \times 10^7$ cells/mL. The cells were encapsulated in MPNs by adding a 10 μL TA solution (80 mg/mL) and a 10 μL $FeCl_3 \cdot 6H_2O$ solution (20 mg/mL) into a 5 mL cell suspension. The cells were then washed twice with 10 mL TAP solutions. After the desired cycles of coating, the cell density was adjusted to $2.5 \times 10^6$ cells/mL for further examinations.

#### 1.4 Chemical and structural characterizations

##### 1.4.1 Confocal laser scanning microscopy (CLSM)

The existence of the MPN layer was confirmed using a fluorescent probe. DyLight 405 NHS ester-conjugated albumin from bovine serum (BSA-DyLight405) was synthesized using the following protocol. 50 μg DyLight™ 405 NHS ester was dissolved in 5 μL DMF, and 210 μg BSA was dissolved in 420 μL sodium borate buffer (0.05 M, pH 8.5). The two solutions were mixed and reacted for 2 h in darkness at room temperature. The product was purified by dialysis in 50 mL 1× PBS for three days in darkness at 4 °C, using a MINI dialysis device (10k MWCO, Slide-A-Lyzer, Thermo Fisher Scientific, MA, USA). The BSA-DyLight405 stock solutions were stored at 4 °C for up to a month.

Inactivated cells for stability tests were prepared by overnight exposure to UV-C radiation, and stained with PI. PI stock solution was prepared by dissolving PI in DI water (1 mg/mL) and stored in darkness at 4 °C.

For fluorescent labeling, the cells in a 200 μL cell suspension were collected by centrifugation (2000 × g, 5 min), washed twice with fresh TAP media, resuspended in a 200 μL BSA-DyLight405 stock solution, and incubated for 15 min in darkness. For stability test, 1 μL PI stock solution was added, and incubated in darkness for 15 min. The labeled cells were then washed twice with fresh TAP media, resuspended in TAP media, and characterized by CLSM (FLUOVIEW FV1200, Olympus, Tokyo, Japan). The BF was chosen for the observation of *C. reinhardtii*, the Alexa Fluor 405 channel for BSA-DyLight405-labeled MPN layers, and the Texas red channels for the autofluorescence of *C. reinhardtii*. For stability test, the Texas red channel was chosen for the observation of inactivated *C. reinhardtii*.

##### 1.4.2 Transmission electron microscopy (TEM)

The morphologies of the microalgae cells were observed by TEM (JEM-1400, JEOL Ltd., Tokyo, Japan). Before adding the samples, the carbon-coated copper grids (200 mesh) were treated by a JEOL HDT-400 hydrophilic treatment device (JEOL Ltd., Tokyo, Japan). A 2 μL suspension of concentrated cells was placed on a hydrophilized copper grid for 2 min. Afterwards, the excess cell suspension was removed by blotting with a filter paper.

##### 1.4.3 Optical microscopy

The morphologies of the microalgae cells were characterized by optical microscopy using an inverted microscope (Eclipse TE2000-U, Nikon, Tokyo, Japan). A cell suspension (10 μL) was gently dropped onto a glass slide, immediately covered with a coverslip, and observed in both BF and a green filtered mode. Digital images were acquired using an integrated CCD camera.

##### 1.4.4 Attenuated total reflectance Fourier transform infrared (ATR-FTIR) spectroscopy

The biomolecular constituents of the microalgae cells were characterized by ATR-FTIR spectroscopy (IMV-4000, JASCO Ltd., Tokyo, Japan). The microalgae cells were washed with DI water, collected by centrifugation (2000 × g, 5 min), dried under vacuum, and examined in 128 scans over 650–4000 $cm^{-1}$ at a resolution of 8.0 $cm^{-1}$.

#### 1.5 Viability and functional health assays

##### 1.5.1 Live assay

The cell viability was investigated by live staining with FDA. FDA stock solution was prepared by dissolving FDA in acetone (5 mg/mL), and stored in darkness at –20 °C. Before the live assay, the cells were washed with and resuspended in a fresh TAP medium. Then, a 4 μL FDA stock solution was added to a 200 μL cell suspension. The cells were stained in darkness for 15 min, collected by centrifugation (2000 × g, 5 min), washed twice with TAP media, resuspended in a 200 μL TAP medium, and observed with a fluorescent microscope (Eclipse TE2000-U, Nikon, Tokyo, Japan). Bright field (BF) and a green filter were chosen to observe *C. reinhardtii* and FDA-stained live cells, respectively. The cells in the BF were counted as the total number of cells, and the cells in the green filter were counted as the number of live cells. At least 200 cells were counted for each sample, and the absolute cell viability and relative cell viability were defined with Eq. S1 and Eq. S2, respectively.

$$Absolute\ cell\ viability = \frac{Number\ of\ live\ cells}{Total\ number\ of\ cells} \times 100\% \qquad \text{(Eq. S1)}$$

where:

$Number\ of\ live\ cells$ was the number of cells observed in the green filter;

and $Total\ number\ of\ cells$ was the number of cells observed in the BF.

$$Relative\ cell\ viability\ at\ t_i = \frac{Absolute\ cell\ viability\ at\ t_i}{Absolute\ cell\ viability\ at\ t_0} \times 100\% \qquad \text{(Eq. S2)}$$

where:

$Relative\ cell\ viability\ at\ t_i$ was determined by normalizing the absolute cell viability at a predetermined time ($t_i$) with the absolute cell viability at the beginning time ($t_0$);

$Absolute\ cell\ viability\ at\ t_0$ was investigated immediately after the MPN encapsulation processes for each measurement;

and $Absolute\ cell\ viability\ at\ t_i$ was investigated at $t_i$.

In the case of heated cells, both the survival rates and the densities of integral cells before and after the thermal treatments were considered for the expression of cell viability (Eq. S3) because of heat-induced lysis (Fig. S12).

$$Relative\ cell\ viability\ at\ t_i = \frac{Survival\ rate\ at\ t_i \times Density\ at\ t_i}{Survival\ rate\ at\ t_0 \times Density\ at\ t_0} \times 100\% \qquad \text{(Eq. S3)}$$

##### 1.5.2 Mitochondrial membrane potential assay

The mitochondrial membrane potential of *C. reinhardtii* was determined using a JC-1 mitochondrial membrane potential (MMP) detection kit, which contained JC-1 dye and an imaging buffer (10×). JC-1 stock solution was prepared by adding 50 μL DMSO into the tube of JC-1 dye (100 nmol), and gently mixing through pipetting. The JC-1 stock solution was stored in darkness at –20 °C for up to one month. The JC-1 working solution was freshly prepared within the day by diluting an 18 μL JC-1 stock solution in a 9 mL TAP medium. The imaging buffer solution was freshly prepared within the day by diluting the imaging buffer (10×) ten times in DI water. For MMP assay, a suspension of *C. reinhardtii* was washed with DI water twice and resuspended in a TAP medium. A 0.5 mL JC-1 working solution was added into a 0.5 mL cell TAP suspension. The mixture was incubated in darkness for 30 min and washed with TAP media twice. The cells were then resuspended in a 1.5 mL imaging buffer solution. The fluorescence of the sample was measured at $\lambda_{ex}$ = 488 nm and at $\lambda_{em}$ = 538 nm or $\lambda_{em}$ = 596 nm (Fig. S4), using a spectrofluorometer (FP-8300, JASCO, Tokyo, Japan). The baseline calibration of the fluorescence emission spectra was conducted using the OriginPro software. The mitochondrial membrane potential was determined by the ratio between the fluorescence intensity at $\lambda_{em}$ = 596 nm and $\lambda_{em}$ = 538 nm (Eq. S4).

$$R = \frac{I_f^{488\to596}}{I_f^{488\to538}} \quad \text{(Eq. S4)}$$

where:
$R$ was the fluorescence intensity index;
$I_f^{488\to596}$ was the fluorescence intensity at $\lambda_{ex}$ = 488 nm and $\lambda_{em}$ = 596 nm;
and $I_f^{488\to538}$ was the fluorescence intensity at $\lambda_{ex}$ = 488 nm and $\lambda_{em}$ = 538 nm.

#### 1.6 Biochemical composition and metabolism assays

##### 1.6.1 Pigment assay

A suspension of *C. reinhardtii* was collected and washed with DI water, and the pigments in the culture were extracted with ethanol in darkness for 1 h. The cell debris was spined down (2000 × g, 20 min), and the absorbance of the supernatant was measured with an ultraviolet–visible (UV–vis) spectrophotometer (Nanodrop™ One, Thermo Fisher Scientific, MA, USA). The concentrations of chlorophyll a (Chl a), chlorophyll b (Chl b), and full carotenoids (Caro) were calculated using the following equations.[1]

$$[Chl\ a] = 13.95A_{665} - 6.88A_{649} \quad \text{(Eq. S5)}$$

$$[Chl\ b] = 24.96A_{649} - 7.32A_{665} \quad \text{(Eq. S6)}$$

$$[Caro] = (1000A_{470} - 2.05[Chlorophyll\ a] - 114.8[Chlorophyll\ b])/245 \quad \text{(Eq. S7)}$$

where:
$[X]$ was the concentration of pigment $X$;
and $A_n$ was the UV–vis absorbance at $n$ nm.

To account for the potential variability in the initial states of cells across independent groups, the metabolite production index was introduced. This index was calculated by normalizing the metabolite concentration at the time point of interest to the concentration measured before any treatment (Eq. S8), allowing for more accurate comparisons across experimental conditions.

$$Index = \frac{C_i/N_i}{C_0/N_0} \quad \text{(Eq. S8)}$$

where:
$Index$ was the metabolite production index of pigments, starches, or lipids;
$C_i$ was the measured concentration of this metabolite at time $t_i$;
$N_i$ was the cell density at time $t_i$;
$C_0$ was the measured concentration of this metabolite of an untreated control;
and $N_0$ was the cell density of this control.

##### 1.6.2 Lipid assay

The lipid production of *C. reinhardtii* was visualized with Nile red. Nile red stock solution was prepared by dissolving Nile red in acetone (1 mg/mL), and stored in darkness at –20 °C. Before the lipid assay, the cells were washed with and resuspended in fresh TAP media. Then, a 1 μL Nile red stock solution was added to a 1 mL cell suspension. The cells were stained in darkness for 10 min, collected by centrifugation (2000 × g, 5 min), washed twice with TAP media, resuspended in a 200 μL TAP medium, and observed with confocal laser scanning microscopy (FLUOVIEW FV1200, Olympus, Tokyo, Japan). Alexa Fluor 568 and Cy5.5 channels were chosen to observe Nile red-stained lipids and *C. reinhardtii*, respectively.

Total lipid contents of C. reinhardtii were extracted according to the Bligh and Dyer method.[2] Briefly, the microalgae cells from a 30 mL cell suspension were collected by centrifugation (2000 × g, 5 min), followed by the addition of a 6 mL methanol/chloroform solution (2:1 v/v). The mixture was incubated for 30 min before sequentially adding 2 mL chloroform and 3.6 mL DI water. After thorough mixing, the sample was centrifugated at 5000 × g for 10 min. The lower phase was then collected, dried via solvent evaporation, and weighed.

##### 1.6.3 Starch assay

The starch production of *C. reinhardtii* was determined using a starch assay kit, which contained starch assay reagent, glucose (HK) assay reagent, and starch assay standard. The starch assay reagent and the glucose assay reagent were reconstituted in 20 mL DI water respectively, mixed several times by inversion, and stored at 4 °C for up to one month.

The samples for starch assay were prepared as follows. The microalgae cells from a 1 mL suspension were collected by centrifugation (2000 × g, 5 min), frozen in liquid nitrogen, and stored at –80 °C. Before starch assay, the cells were thawed in 1 mL methanol at –20 °C for 1 h. The pellet was collected through centrifugation (18000 × g, 15 min), washed with 1 mL sodium acetate solution (1 M), and suspended in 1 mL DI water. The mixture was sonicated at 40 °C for at least 1 h until no visible solids remained in suspension. The samples were then autoclaved at 121 °C under standard pressure (approximately 0.1 MPa) for 1 h and subsequently maintained at 60 °C.

Test tubes for starch assay were prepared as described in Table S9, mixed well, incubated at 60 °C for 15 min, and cooled to room temperature.

Test tubes for glucose assay were prepared as described in Table S10, mixed well, incubated at room temperature for 15 min, and the absorbance was measure at 340 nm with a UV–vis spectrophotometer.

The absorbance contributed from the blanks was calculated with Eq. S9. The actual absorbance of the tested sample was calculated with Eq. S10. The starch concentration in the prepared samples was calculated with Eq. S11.

$$A_{\text{Total Blank}} = A_{\text{Sample Blank}} - A_{\text{Glucose Assay Reagent Blank}} + A_{\text{Starch Assay Reagent Blank}} \quad \text{(Eq. S9)}$$

$$\Delta A = A_{\text{Sample for Test}} - A_{\text{Total Blank}} \quad \text{(Eq. S10)}$$

where:

$\Delta A$ was the actual absorbance of the tested sample;

and $A_X$ was the absorbance of test tube $X$ measured at 340 nm.

$$SC = \frac{(\Delta A)\left(\frac{TVSA}{SVSA}\right)\left(\frac{TVGA}{SVGA}\right)(\text{Starch } MW)}{(\varepsilon)(d)} \quad \text{(Eq. S11)}$$

where:

$SC$ was starch concentration in μg/mL;

$TVSA$ was total assay volume from starch assay in mL, and $TVSA = 1$ mL;

$SVSA$ was sample volume from starch assay in mL, and $SVSA = 0.5$ mL;

$TVGA$ was total assay volume from glucose assay in mL, and $TVGA = 1$ mL;

$SVGA$ was sample volume from glucose assay in mL, and $SVGA = 0.5$ mL;

$Starch\ MW$ was the molecular weight of starch, and Starch $MW = 162.1$ g/mol;

$\varepsilon$ was millimolar extinction coefficient for nicotinamide adenine dinucleotide (NADH) at 340 nm, and $\varepsilon = 6.22$ L/mmol/cm;

$d$ was the light path, and $d = 1$ cm.

#### 1.7 Quantitative and growth measurements

##### 1.7.1 Cell density measurement

The cell density of a *C. reinhardtii* suspension was measured using a standard hemocytometer (Thoma, Hirschmann, Germany) under a light microscope (Eclipse TE2000, Nikon, Tokyo, Japan).

##### 1.7.2 Cell growth curve measurement

A suspension of *C. reinhardtii* was diluted in a 40 mL TAP medium with its density adjusted to $1\times10^5$ cells/mL. The cells were cultured in an LED-equipped shaking incubator (LC-LED450W and BR-43FL·MR, Taitec, Saitama, Japan) at 25 °C at 120 rpm under continuous light, continuous dark, or 8 h/16 h light/dark cycles. The light intensity was set to approximately 120 μmol photons $m^{-2}$ $s^{-1}$. The cell density was measured every 24 h. The growth curve fitting was performed with OriginPro software by using S logistic fitting function. The growth rate $\mu$ and doubling time $t_d$ was calculated with Eq. S12 and Eq. S13.

$$\mu = \frac{\ln N_2 - \ln N_1}{t_2 - t_1} \quad \text{(Eq. S12)}$$

$$t_d = \frac{\ln 2}{\mu} \quad \text{(Eq. S13)}$$

where:

$\mu$ was the growth rate;

$t_d$ was the growth rate;

and $\ln N_i$ was the natural logarithm of cell density at time $t_i$.

Since it is difficult to precisely identify the time duration of the lag phases, the delay in cell growth was quantified as the difference in time to reach half of the maximum cell density ($\Delta T_{50}$) between algae@MPN and bare algae.

### 2. Supplementary figures

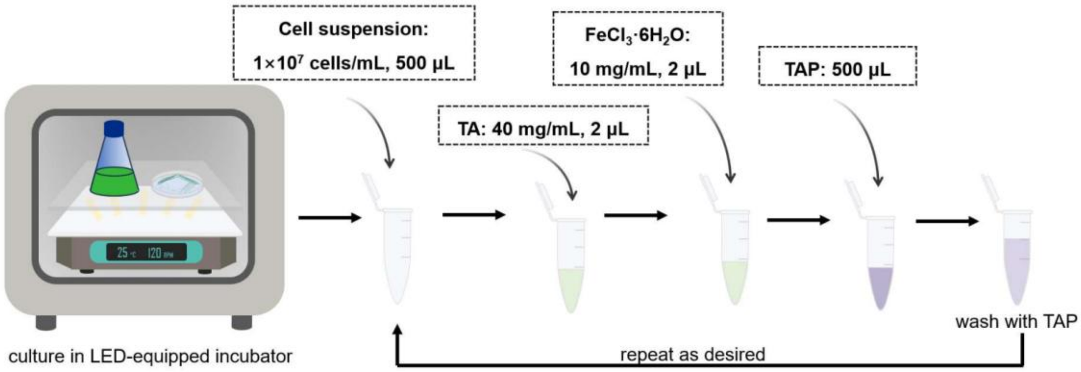

**Figure S1.** The process of encapsulating *C. reinhardtii* microalgae cells into MPN nanofilms.

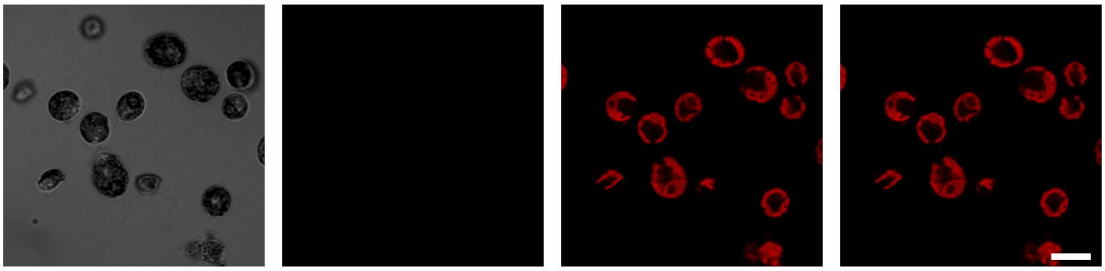

**Figure S2.** CLSM images of bare algae labeled with BSA-DyLight405. Scale bar: 10 μm.

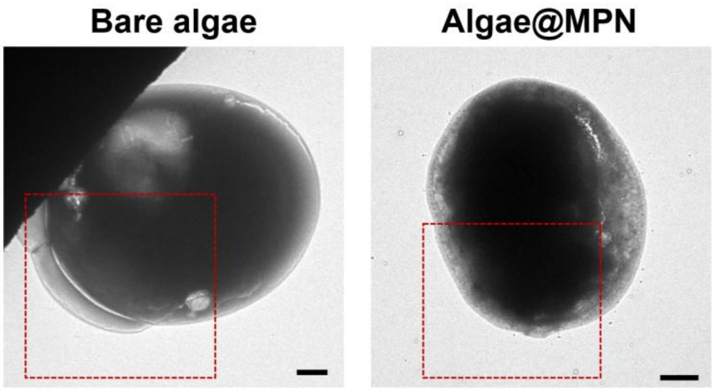

**Figure S3.** TEM images of bare algae and algae@MPN. Scale bars: 2 μm.

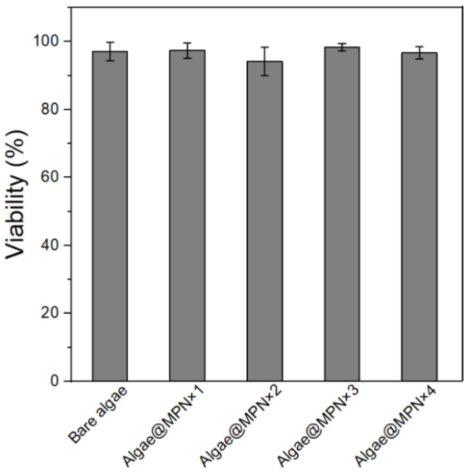

**Figure S4.** Viability of bare algae and algae@MPN quantified by FDA staining. Error bars represent standard deviations (n = 3).

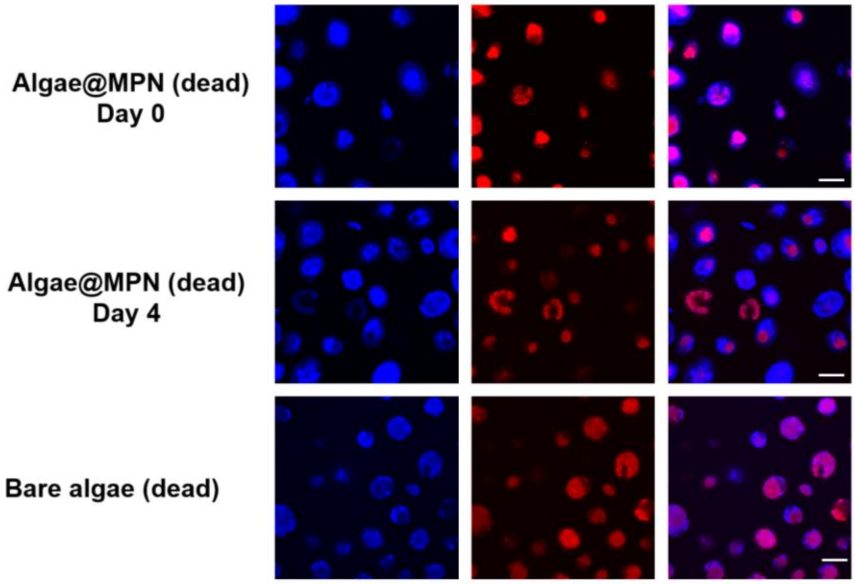

**Figure S5.** CLSM images of inactivated bare algae and algae@MPN stained with BSA-DyLight405 and PI to highlight that the MPN shell survived 4 days in cell culture media. Scale bars: 10 μm.

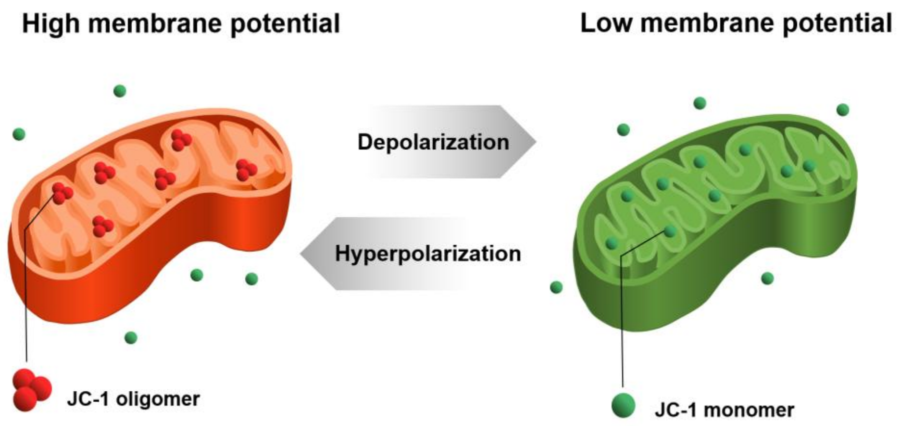

**Figure S6.** JC-1-based mitochondrial membrane potential detection. Under high MMP of healthy mitochondria, JC-1 concentrates and forms oligomers that emit red fluorescence; under low MMP of depolarized mitochondria during apoptosis or other dysfunctions, JC-1 remains in its monomeric form and emits green fluorescence.

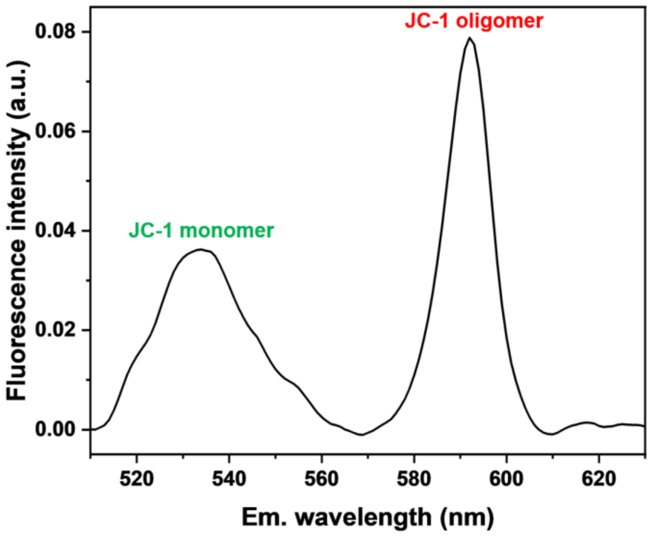

**Figure S7.** Emission spectrum of JC-1-dyed *C. reinhardtii* under 488 nm excitation.

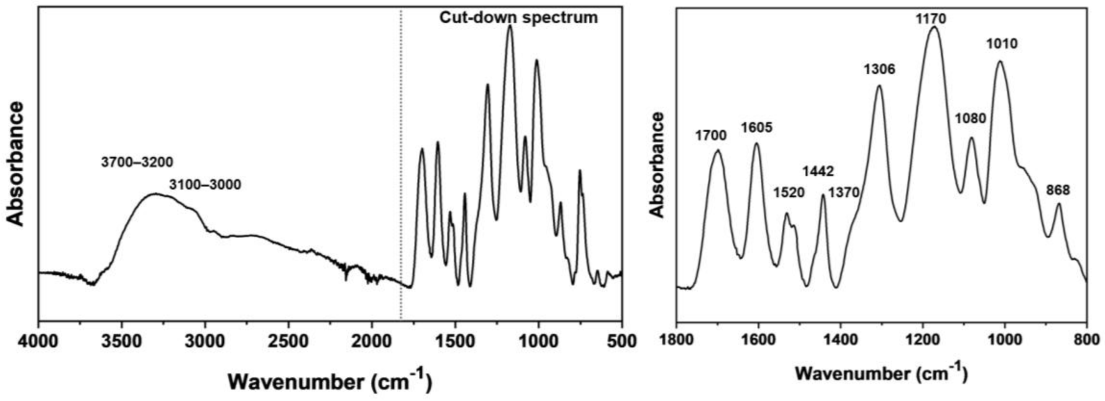

**Figure S8.** ATR-FTIR spectrum and cut-down spectrum of 1800–500 $cm^{-1}$ of TA.

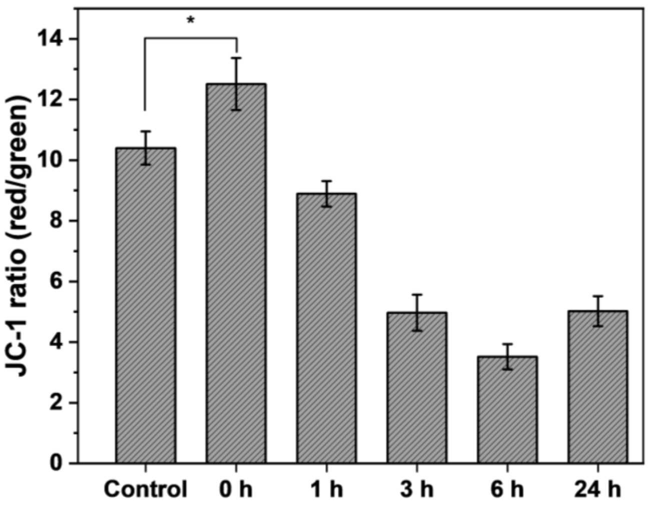

**Figure S9.** The red-to-green fluorescence ratios of algae@MPN incubated in darkness for various times and measured with JC-1 MMP assay. Control: untreated bare algae. Error bars represent standard deviations of n = 3 replicates. Statistical analysis was performed using one-way *ANOVA* (*$p < 0.05$).

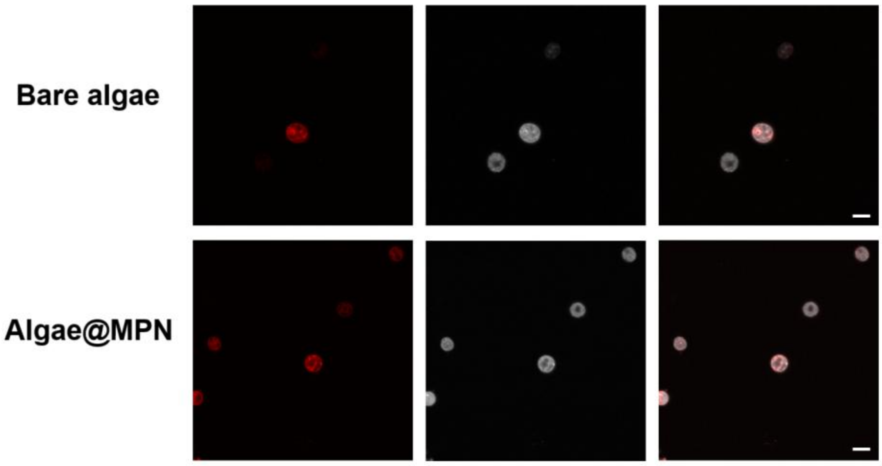

**Figure S10.** CLSM images of Nile red-stained bare algae and algae@MPN after dark cultivation for 7 days. Lipid droplets (left) and chlorophyll autofluorescence (middle) are shown, together with merged images (right). Independent experiments were performed (n = 3) with similar results. Scale bars: 10 μm.

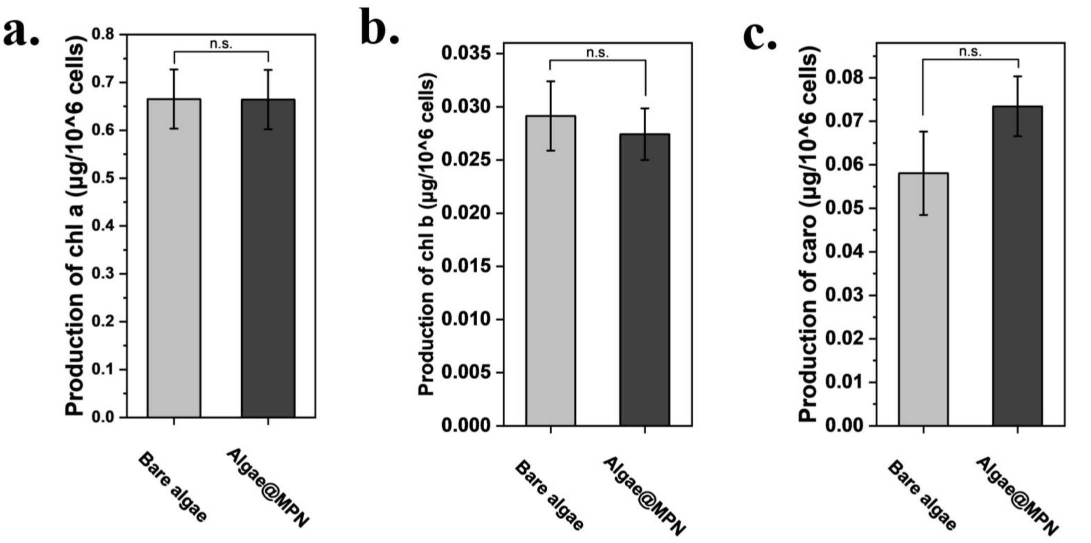

**Figure S11.** Long-term accumulation of (a) chlorophyll a (Chl A), (b) chlorophyll b (Chl B), and (c) total carotenoids (Caro) of bare algae and algae@MPN for 4 days under light. Error bars represent standard deviations (n = 3). Statistical analysis was performed using one-way *ANOVA* (*n. s. p* $\geqslant$ 0.05).

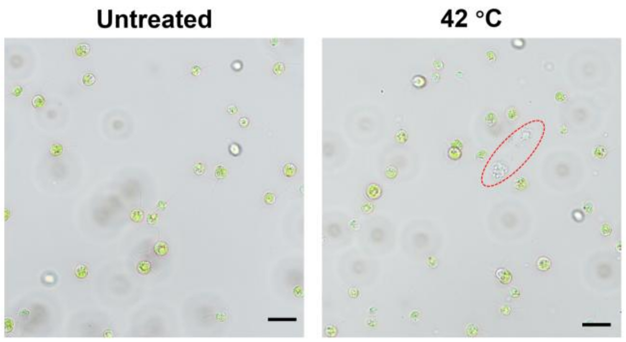

**Figure S12.** Optical microscope images of bare algae untreated, and treated at 42 °C for 30 min. The red circled regions are the debris of dead cells. Scale bars: 2 μm.

#### 3. Supplementary tables

**Table S1.** ATR-FTIR peak assignments for microalgal cells.

| Peak type | Wavenumber ($cm^{-1}$) | Assignment |
|---|---|---|
| Mutual peaks | 1740 | ν(C=O) of lipids and fatty acids |
| | 1643 | ν(C–O) of proteins (amide I) |
| | 1540 | δ(N–H) and ν(C–N) of proteins (amide II) |
| | 1454 | δ($CH_2$) and δ($CH_3$) of lipids and proteins |
| | 1404 | δ(C–H) of methylene |
| | 1240 | ν(P=O) of phosphodiester in nucleic acids and phospholipids |
| | 1200–900 | ν(C–O–C) of polysaccharides |
| | 876 | ν(P–O–C) of phosphodiester in nucleic acids, and δ(C–H) of aromatic amino acids |
| Unique to algae@MPN | 1439 | δ(O–H) of carboxylic acids of TA |
| | 1344 | ν(C–O) of carboxylic acids and esters of TA |
| | 1311 | δ(O–H) of phenols of TA |
| | 1204 | ν(C–O) of esters of TA |

ν, stretching vibration; δ, bending vibration.

**Table S2.** ATR-FTIR peak assignments for TA.

| Wavenumber ($cm^{-1}$) | Assignment |
|---|---|
| 3700–3200 | ν(O–H) of phenol groups and water |
| 3100–3000 | ν(C–H) of aromatic rings |
| 1700 | ν(C=O) of ester groups |
| 1605 | ν(C=C) of aromatic rings |
| 1520 | ν(C=C) of aromatic rings |
| 1442 | δ(O–H) of carboxylic acid |
| 1370 | ν(C–O) of carboxylic acid |
| 1306 | δ(O–H) of phenol groups |
| 1170 | ν(C–O) of ester groups |
| 1080 | ν(C–O) of ester groups |
| 1010 | ν(C–C) and δ(C–O) of aromatic rings |
| 868 | δ(C–H) of aromatic rings |

ν, stretching vibration; δ, bending vibration.

**Table S3.** Normalized peak areas of ATR-FTIR spectra for microalgae.

| Peak center | Bare algae | $H_2O_2$-treated bare algae | $H_2O_2$-treated algae@MPN | Relevant biomolecules |
|---|---|---|---|---|
| 1740 $cm^{-1}$ | $1.5\times10^{-2}$ | $2.5\times10^{-2}$ (↑↑) | $1.8\times10^{-2}$ (–) | Lipids |
| 1540 $cm^{-1}$ | $4.4\times10^{-1}$ | $5.0\times10^{-1}$ (–) | $6.1\times10^{-1}$ (↑) | Proteins |
| 1240 $cm^{-1}$ | $2.1\times10^{-1}$ | $5.5\times10^{-1}$ (↑↑↑) | $1.1\times10^{-1}$ (↓) | Nucleic acids, and phospholipids |
| 876 $cm^{-1}$ | $3.8\times10^{-2}$ | $2.3\times10^{-1}$ (↑↑↑) | $6.0\times10^{-3}$ (↓↓) | Nucleic acids, proteins, and sugars |

The amide I band at 1643 $cm^{-1}$ was used as the reference for normalization. –: no significant change (change magnitude < ±20%); ↑ or ↓: modest change (±20% to ±50%); ↑↑ or ↓↓: substantial change (> ±50%); ↑↑↑ or ↓↓↓: significant change (> ±100%). The change magnitude was calculated by comparing each value to the bare algae sample as a control, using the Eq. S14. Positive value indicates an increase, and negative value indicates a decrease relative to the Control.

$$\text{Change magnitude (\%)} = \frac{\text{Sample} - \text{Control}}{\text{Control}} \times 100\% \qquad \text{(Eq. S14)}$$

**Table S4.** Growth kinetics data obtained from the microalgae growth curves.

| Light condition | Sample | $N_{max}$ (cells/mL) | $\mu$ ($h^{-1}$) | $t_d$ (h) | $T_{50}$ (h) | $\Delta T_{50}$ (h) |
|---|---|---|---|---|---|---|
| Continuous light | Bare algae | $1.73\times10^{7}$ | 0.13 | 5.3 | 43.3 | - |
| | Algae@MPN | $1.62\times10^{7}$ | 0.20 | 3.5 | 48.9 | 5.6 |
| Light/dark cycle | Bare algae | $9.57\times10^{6}$ | 0.045 | 15.4 | 67.3 | - |
| | Algae@MPN | $8.59\times10^{6}$ | 0.046 | 15.1 | 76.2 | 8.9 |
| Continuous dark | Bare algae | $3.18\times10^{6}$ | 0.017 | 40.8 | 114.8 | - |
| | Algae@MPN | $3.07\times10^{6}$ | 0.021 | 33.0 | 126.7 | 11.9 |

growth rates ($\mu$), doubling times ($t_d$)

**Table S5.** Recipe for TAP medium.

| Component | TAP broth | TAP agar |
|---|---|---|
| TAP-salt solution | 10 mL | 10 mL |
| Phosphate solution | 1 mL | 1 mL |
| Trace elements solution | 10 mL | 10 mL |
| Agar | - | 15 g |
| DI water | 1 L | 1 L |

**Table S6.** Recipe for TAP-salt solution.

| **Component** | **Amount** |
|---|---|
| $NH_4Cl$ | 40.0 g |
| $MgSO_4·7H_2O$ | 10.0 g |
| $CaCl_2·2H_2O$ | 5.0 g |
| DI water | 1 L |

**Table S7.** Recipe for phosphate solution.

| **Component** | **Amount** |
|---|---|
| $K_2HPO_4$ | 10.8 g |
| $KH_2PO_4$ | 5.6 g |
| DI water | 1 L |

**Table S8.** Recipe for trace elements solution.

| **Component** | **Amount** |
|---|---|
| $Na_2·EDTA$ | 5.0 g |
| $ZnSO_4·7H_2O$ | 2.2 g |
| $H_3BO_3$ | 1.14 g |
| $MnCl_2·4H_2O$ | 0.51 g |
| $CoCl_2·6H_2O$ | 0.16 g |
| $CuSO_4·5H_2O$ | 0.16 g |
| $(NH_4)_6Mo_7O_{24}·4H_2O$ | 0.11 g |
| $FeSO_4·7H_2O$ | 0.50 g |
| KOH | 1.6 g |
| DI water | 1 L |

**Table S9.** Recipe for the test tubes for starch assay.

| **Test tube** | **Starch assay reagent (mL)** | **Sample (mL)** | **DI water (mL)** |
|---|---|---|---|
| Starch assay reagent blank | 0.5 | - | 0.5 |
| Glucose assay reagent blank | - | - | 1 |
| Sample blank | - | 0.5 | 0.5 |
| Sample for test | 0.5 | 0.5 | - |

**Table S10.** Recipe for the test tubes for glucose assay.

| Test tube | Glucose assay reagent (mL) | Sample from starch assay (mL) |
|---|---|---|
| Starch assay reagent blank | 0.5 | 0.5 |
| Glucose assay reagent blank | 0.5 | 0.5 |
| Sample blank | 0.5 | 0.5 |
| Sample for test | 0.5 | 0.5 |
